## Supplementary Table 1 for "Human DDX6 regulates translation and decay of inefficiently translated mRNAs"

**Supplementary Table 1. Constructs and mutants used in this study.**

<table>
<tr><th>Name</th><th>Fragments / mutations</th><th>Plasmid</th></tr>
<tr><td rowspan="2">R-LUC</td><td>-</td><td>pCIneo-RLuc</td></tr>
<tr><td colspan="2"><u>Last 30 codon sequence</u><br>AAG GGC CTC CAC TTC AGC CAG GAG GAC GCT<br>CCA GAT GAA ATG GGT AAG TAC ATC AAG AGC<br>TTC GTG GAG CGC GTG CTG AAG AAC GAG CAG</td></tr>
<tr><td rowspan="2">R-LUC 30xRC</td><td>30xRC</td><td>pCIneo-RLuc_30xRC</td></tr>
<tr><td colspan="2"><u>Last 30 codon sequence</u><br>AAA GGT CTA CAT TTT TCG CAA GAA GAT GCG<br>CCG GAT GAA ATG GGT AAA TAT ATA AAA TCG<br>TTT GTA GAA CGT GTA CTA AAA AAT GAA CAA</td></tr>
<tr><td>MBP-Strep</td><td>-</td><td>pnEK-NvHM-Strep-MBP</td></tr>
<tr><td>NusA-Strep-DDX6</td><td>-</td><td>pETM-60-NusA-3C-HsRCK_296-472-Strep</td></tr>
<tr><td>GFP-MBP</td><td>-</td><td>pT7-EGFP-C1-MBP</td></tr>
<tr><td rowspan="6">GFP-DDX6</td><td>-</td><td>pT7-EGFP-C1-HsDDX6</td></tr>
<tr><td>N-ter</td><td>pT7-EGFP-C1-HsDDX6_1-295</td></tr>
<tr><td>C-ter</td><td>pT7-EGFP-C1-HsDDX6_296-463</td></tr>
<tr><td>E236Q</td><td>pT7-EGFP-C1-HsDDX6_E236Q</td></tr>
<tr><td>Mut1</td><td>pT7-EGFP-C1-HsDDX6_Mut1</td></tr>
<tr><td>Mut2</td><td>pT7-EGFP-C1-HsDDX6_Mut2</td></tr>
<tr><td>HA-RPL22</td><td>-</td><td>pCIneo-HA-RPL22</td></tr>
<tr><td>RL-AR</td><td>-</td><td>pCIneo-RL-AR</td></tr>
<tr><td>RL-stop-AR</td><td>-</td><td>pCIneo-RL-stop-AR</td></tr>
<tr><td>RL-BMP2</td><td>-</td><td>pCIneo-RL-BMP2</td></tr>
<tr><td>RL-stop-BMP2</td><td>-</td><td>pCIneo-RL-stop-BMP2</td></tr>
<tr><td>V5-SBP-MBP-MS2</td><td>-</td><td>pCIneo-v5-SBP-MBP-MS2</td></tr>
<tr><td>RL-6xMS2bs</td><td>-</td><td>pCIneo-RL-6xMS2bs</td></tr>
<tr><td>RL-AR-6xMS2bs</td><td>-</td><td>pCIneo-RL-AR-6xMS2bs</td></tr>
<tr><td>RL-BMP2-6xMS2bs</td><td>-</td><td>pCIneo-RL-BMP2-6xMS2bs</td></tr>
</table>

### SUPPLEMENTAL FIGURES

**Figure S1 Characterization of HEK293T DDX6 KO cells.**

**A.** Immunoblots were probed with antibodies recognizing DDX6 and Tubulin.

**B, C.** Multidimensional scaling (MDS) analysis for the Ribo-Seq (B) and RNA-Seq (C) replicate libraries from HEK293T wild-type (WT) and DDX6 KO cells. The Ribo-Seq and RNA-Seq experiments were reproducible as replicates clustered together.

**D.** Sanger sequencing of the DDX6 genomic region targeted by the DDX6 sgRNA. Frameshift mutations were detected in exon 3 of both alleles. These generate premature STOP codons (PTC) and deletions in DDX6.

**E.** Northern blot analysis of CNOT3 tethered to an R-LUC reporter mRNA in HEK293T WT or DDX6 KO cells. Indicated cells were transfected with a mixture of three plasmids: 1. expressing the Renilla luciferase (R-LUC) containing 5BoxB reporter, 2. expressing the Firefly luciferase (F-LUC) as a transfection control, and 3. expressing the λN-HA peptide (−) or λN-HA-CNOT3 (+).

**F.** Ribosome footprints (RFP) and total mRNA (RNA) reads distribution along *DDX6* mRNA in wild-type (WT) and DDX6 KO cells. Of note, RFP and total RNA counts for DDX6 are drastically reduced in the KO cells.

**Figure S2 Identification of DDX6 target mRNAs.**

**A.** Schematic representation of the experimental strategy to identify mRNAs targeted by DDX6 for translational repression and decay.

**B-G.** Ribosome footprints (RFP) and total mRNA (RNA) reads distribution along DDX6 target mRNAs in wild-type (WT) and DDX6 KO cells. Potential ribosome stalling sites are indicated in red dotted boxes.

**Figure S3 Characterization of DDX6 target mRNAs.**

**A.** qPCR analysis of *LGALS1*, *DLX5*, *ENO2* and *PSMB9* mRNA levels in HEK293T wild-type (WT) and DDX6 KO and rescued with GFP-tagged DDX6 (wild-type or the indicated mutants). $log_2$FC values for each transcript as determined by the RNA-seq experiments are indicated.

**B-D.** Ridgeline plots illustrating the GC content (B), coding sequence (CDS) length (C), and translational efficiency (TE) (E) of translationally regulated DDX6 target mRNAs. Statistical significance was calculated with the one-sided Wilcoxon rank sum test.
